# Adult-type diffuse gliomas share recurring cell states driven by a common astrocyte-like glioma stem cell population

**DOI:** 10.1101/2024.06.13.598923

**Authors:** Walter Muskovic, Ashraf Zaman, Madhavi Pandya, Emily Holton, Chia-Ling Chan, Rachael A McCloy, Himanshi Arora, Venessa T Chin, Joseph E Powell

**Affiliations:** Translational Genomics, Garvan Institute of Medical Research, The Kinghorn Cancer Centre, Darlinghurst, NSW 2010, Australia; School of Clinical Medicine, The University of New South Wales, Sydney, NSW 2052, Australia; The Kinghorn Cancer Centre, St Vincent’s Hospital Sydney, Darlinghurst, NSW 2010, Australia; UNSW Cellular Genomics Futures Institute, University of New South Wales, Sydney, NSW 2052, Australia

**Author notes:** co-first (Walter Muskovic, Ashraf Zaman). Walter Muskovic –, Ashraf Zaman –, Madhavi Pandya –, Emily Holton –, Chia-Ling Chan –, Rachael A McCloy –, Himanshi Arora –, Venessa T Chin –, Joseph E Powell –.

**Keywords:** Glioma, single-cell genomics, glioma stem cell, glioma cell states, glioblastoma, astrocytoma, oligodendroglioma

## Abstract

Adult-type diffuse gliomas are a family of aggressive brain tumours with few effective treatments. Their complex cellular makeup adds to the challenge of finding successful therapies. This intratumoural heterogeneity is fuelled by a subpopulation of glioma stem-like cells (GSCs) that drive tumour growth and resistance to standard treatments. Previous research focused on the three glioma types (astrocytoma, oligodendroglioma, glioblastoma) individually revealed malignant cells mimic the transcriptional profiles of normal brain cell types. Whether these diverse cellular states stem from a shared biological origin is unknown. Here, we show through single-cell RNA sequencing of 40 glioma tumours that all gliomas are described by seven recurring cell states. We also identify a shared astrocyte-like GSC population. Our unique method of identifying GSCs, based on reconstructed tumour phylogenies, repositions astrocyte-like cells at the apex of a differentiation hierarchy in glioma. Our findings indicate the transcriptional heterogeneity observed in gliomas stems from a GSC population recapitulating lineages of healthy adult neural stem cells. These results suggest a shared lineage drives the intratumoural heterogeneity observed in adult-type diffuse gliomas. We anticipate that a deeper understanding of the molecular mechanisms maintaining the GSC state will provide a new framework for future therapeutic development and research into glioma cell biology.

**Highlights:**

- Recurring cell states are shared across adult-type diffuse gliomas
- Reconstructed tumour phylogenies identify an astrocyte-like glioma stem cell population
- Tumour subclones are segregated non-randomly across cell states

## Introduction

Adult-type diffuse gliomas collectively account for most adult neuro-oncology cases and remain incurable due to their intrinsic tumour heterogeneity and the persistence of glioma stem cells (GSCs)^1^. The three tumour types, astrocytoma (IDH-mutant), oligodendroglioma (IDH-mutant and 1p/19q co-deleted), and glioblastoma (IDH-wildtype) differ in their genetic architecture, mitotic activity, histological features, preferential localisation, patient demographics, and clinical outcomes. Despite these differences, morphological examination and, more recently, single-cell RNA-sequencing (scRNA-seq) analysis, indicate malignant cells of all three tumour types mimic cells of astrocytic, oligodendroglial, and neuronal lineages^2–8^. These transcriptional signatures persist despite the unique genetic architecture of each tumour, suggesting a potential common origin in a multipotent GSC. GSCs have been best characterised in glioblastoma, demonstrating potent tumour-initiating ability, self-renewal capacity, and resistance to standard therapies^9,10^. An improved understanding of commonalities in GSCs across the gliomas could reveal shared mechanisms of tumour growth and resistance to standard therapies. This knowledge would provide a framework for developing pan-glioma therapies that target these vulnerabilities, such as selectively eliminating GSCs, targeting the molecular pathways that maintain their quiescence or promote their differentiation.

Previous research, focused individually on glioblastoma, oligodendroglioma, and astrocytoma, has identified glioma cells recapitulate specific brain cell lineages^2–8^. This work hints at a shared origin of GSC-driven intratumoural heterogeneity across gliomas. However, a detailed understanding of whether GSCs represent a truly shared population across these tumour types, and their role in shaping the cellular composition of each tumour type, is lacking. Here, we address this gap with a cohort of glioma tissue samples that include each of the three cancer types. Profiling these samples with scRNA-seq, we employ a novel method to classify multi-dimensional cell states, and identify seven cell states shared amongst gliomas, although differing in their proportions. We identify a prospective common GSC cell state, and validate its identification based on the accumulation of somatic mutations and clonal evolution. This method leverages the genetic relationship between tumour cells and how genetic information flows within a cell hierarchy, originating with GSCs. We inferred copy number variations (CNVs) from the scRNA-seq data to construct clonal phylogenies for each tumour. Since tumour cells pass their CNVs down to their differentiating descendants, later-arising alterations won’t be present in earlier populations of cells, allowing us to identify the progenitor population. We apply this approach to identify a quiescent, astrocyte-like GSC population shared across the three tumour types. To rigorously validate our findings, the reconstructed tumour phylogenies and linkage to GSCs were confirmed through both single-cell DNA-sequencing and analysis of mitochondrial DNA mutations. These findings offer a new perspective on the genomic heterogeneity of gliomas by framing it within the context of normal adult neural stem cell (NSC) differentiation. Our work marks a significant step forward by uncovering shared biological features among adult-type gliomas, thereby suggesting new potential avenues for therapeutic intervention.

## Results

### scRNA-seq captures shared transcriptional heterogeneity in glioma tumour cells

To examine the shared biology amongst the three types of adult-type diffuse gliomas, fresh tumour tissue from seven oligodendrogliomas, nine astrocytomas and twenty-four glioblastomas was collected, dissociated directly following surgery and initially profiled with single-cell RNA-sequencing (scRNA-seq) (**Table S1**). From these samples, post quality control (**Methods**), the transcriptional profiles of 543,088 cells were obtained. Cell types were annotated using marker gene sets (**Figure S1A, Table S2**) and were comprised predominantly of four major cell types: tumour cells, myeloid cells, T cells and oligodendrocytes (**Figure 1A**) as well as a small number of other cell types (**Figure S1B**, **Table S3**). Interestingly, the proportion of non-tumour cells was markedly higher in glioblastoma tissue compared with the other tumour types (**Figure 1B, Table S4**). This may be due to this tumour’s highly infiltrative growth characteristics, resulting in a higher average proportion of non-tumour brain tissue in these specimens. Of the 543,088 total cells, 77,250 oligodendroglioma, 111,063 astrocytoma, and 129,292 glioblastoma tumour cells were identified. To ensure tumour cells were accurately distinguished from normal diploid cells, CNVs were inferred from the scRNA-seq data by measuring shifts in transcript abundance and allelic imbalance in expressed heterozygous single-nucleotide polymorphisms (SNPs)^11^. CNVs were detected in all tumour cells (**Figure 1C**). Reassuringly, hallmark genomic alterations were consistently identified, including 1p/19q co-deletion in oligodendroglioma tumour cells and trisomy seven and monosomy ten in glioblastoma tumour cells, consistent with observations from bulk tumour tissue (**Figure 1D**). Following the identification of tumour cells, gene expression data from all samples was integrated. After data pre-processing, dimension reduction and clustering, cells from all three tumour types separated into distinct expression-based clusters rather than by tumour type (**Figure 1E**).

**Figure 1.**
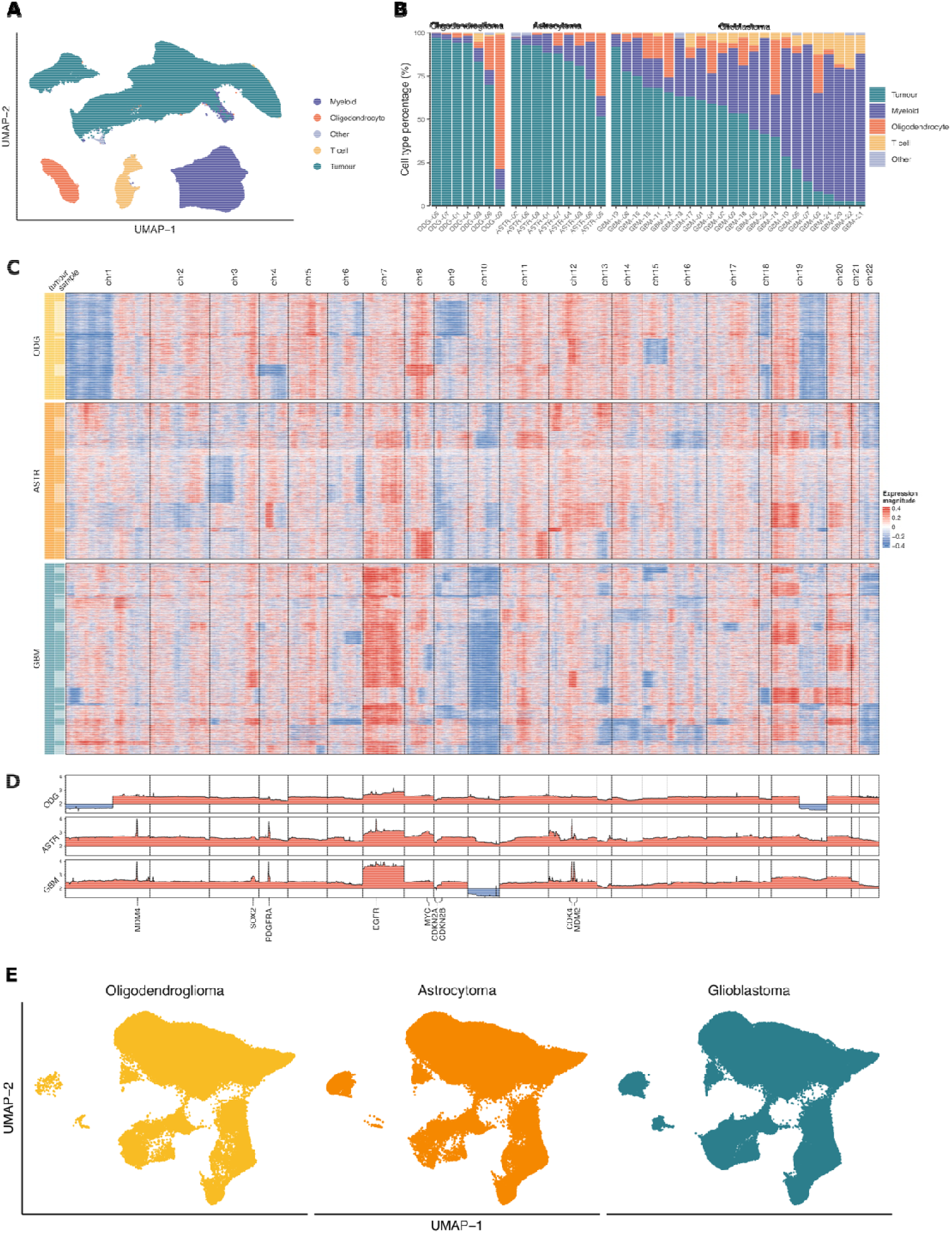
Classification of single cells from 40 adult-type diffuse gliomas. (A) Two-dimensional uniform manifold approximation and progression (UMAP) embedding plot of normal and tumour cells from adult-type diffuse glioma tissue samples. Colours correspond to distinct cell types. Minor cell type populations are grouped as ‘other’ (See Figure S1 for full detail). (B) The percentage of cell types comprising each tumour. Samples are organised left to right by the three tumour types; oligodendroglioma, astrocytoma and glioblastoma. (C) Single-cell heatmap of window-smoothed expression signals across 100 genes. Colours indicate shifts in gene expression relative to cells without CNVs, illustrating inferred copy number events for each chromosome (column). Rows, corresponding to individual cells, are grouped by tumour sample and organised by tumour type: oligodendroglioma (top), astrocytoma (middle) and glioblastoma (bottom). (D) Line plots illustrating the mean gene-level CNV signal across 197 oligodendrogliomas (top), 195 astrocytomas (middle) and 488 glioblastomas (bottom) from The Cancer Genome Atlas (TCGA). The chromosomal location of genes associated with recurring focal CNVs are labelled below. (E) UMAP embedding plot of tumour cells from all samples, merged into one dataset and split by tumour type.

### scRNA-seq identifies seven recurring glioma cell states

To provide a framework for understanding patterns of intratumoural heterogeneity that span tumour types, we sought to define cell states shared across tumours. The high-dimensional gene expression matrix comprised of all tumour cells was decomposed into a small set of underlying factors with non-negative matrix factorisation (NMF) to achieve this aim. An advantage to NMF is that it allows for additive combinations of intrinsic features. For example, a cell expressing genes consistent with an astrocyte-like cell identity may also express genes associated with transient cellular processes such as cell division. The combination of two states defines such a cell. From this analysis we identify seven recurring cell states (**Figure 2A**, **Table S5**) observed in all tumour types. Interestingly, most tumour cells across the three tumour types exist predominantly in a single cell state (**Figure 2A**, **B and Table S6**). This suggests that differentiation pathways remain relatively constrained, with tumour cells committed to a specific lineage rather than a high degree of plasticity or residing in multiple intermediate states.

**Figure 2.**
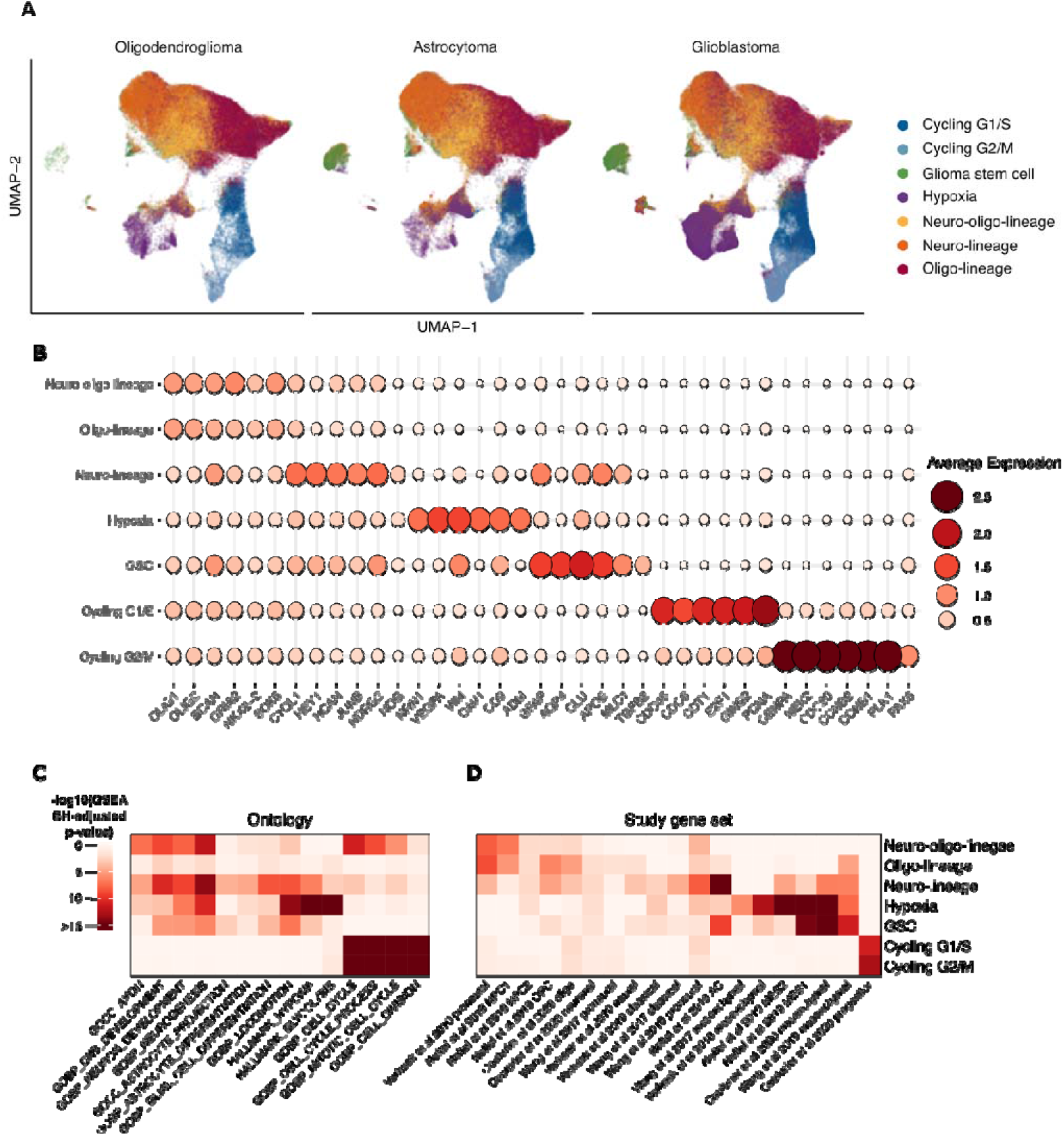
Adult-type diffuse gliomas are defined by seven recurring cell states characterised by distinct cell markers and functional enrichment. (align panels) (A) UMAP embedding plots show tumour cells from each tumour type. Cells are coloured by one of seven recurring transcriptional cell states. (B) Key gene markers define seven distinct cell states. (C) Heatmap of functional enrichment and overlap of the seven identified glioma cell states with cell states identified in existing studies. Columns represent molecular signatures associated with biological processes and states (left) or published cell state gene signatures (right); rows are glioma cell states. Colours represent Benjamini-Hochberg adjusted gene set enrichment analysis *p*-values.

Three cell states showed enrichment of gene ontology terms associated with both oligodendroglial and neuronal lineage development (**Figure 2C**). Of these, cells in one state expressed markers of neural progenitor cells (*HEY1* and *NDRG2*) as well as regulators of neuron growth and adhesion (*C1QL1*, *NCAN* and *SPARCL1*) (herein called neuro-lineage) (**Figure 2B and S2B**). Another state was enriched for expression of key oligodendroglial lineage regulators (*MYRF* and *GPR17*) and oligodendrocyte identity marker genes (*MOG*, *MAG* and *CLDN11*) (herein called oligo-lineage) (**Figure S2D**). A third, intermediate state, which we termed ‘neuro-oligo-lineage’ was identified to be strongly associated with key oligodendrocyte progenitor markers (*OLIG1* and *OLIG2*) as well as neuronal lineage markers (*SOX4*, *SOX11* and *DCX*) (**Figure S2C**).

Two cell states demonstrated high expression of hallmark cell cycle genes and enrichment of associated gene sets (**Figure 2B and 2C**). Expression of genes essential for the initiation and progression of DNA replication (*CENPA*, *NEK2* and *PLK1*) identified the first cell state as capturing cells in the G1/S cell cycle phase. Enrichment of genes controlling mitotic spindle formation and chromosome separation (*CENPA*, *NEK2* and *CDC20*) marked cells in the second state as transitioning through G2/M. These states were marked ‘cycling G1/S’ and ‘cycling G2/M’, respectively. Interestingly, proliferating cells comprised a transcriptionally distinct population that did not overlap other cell states. This suggests that rather than representing a transient state within a differentiating population, the proliferative states capture a distinct cellular program driven by specific signals or transitions within the GSC hierarchy.

A sixth cell state demonstrated strong enrichment of genes and pathways associated with cellular adaptation to compromised oxygen availability, increased glycolysis and promotion of angiogenesis (**Figure 2B and 2C**). Enriched expression of hypoxia-inducible target genes (*EGLN3*, *VEGFA* and *CAV1*) and genes associated with increased glycolysis (*PGK1*, *ENO1* and *ENO2*) imply this state captured a cellular response to low tumour oxygenation and was labelled ‘hypoxia’ accordingly.

Finally, a transcriptionally distinct state was identified that was associated with elevated expression of key astrocyte lineage markers (*GFAP*, *AQP4* and *MLC1*) (**Figure 2B**). These astrocyte-like cells were also enriched for expression of GSC markers *ITGB4* and *S100A4* (**Figure S2F**). *ITGB4* expression levels have recently been shown to correlate with glioma grade and promote GSC self-renewal and gliomagenesis^12^. In addition, *S100A4* has been identified as a key regulator of quiescent GSCs and its expression is correlated with significantly worse prognosis among glioma patients^13^. These observations suggest this astrocyte-like cell state may represent a GSC population and was labelled ‘glioma stem cell’ accordingly.

Having identified these states, we sought to understand how they aligned with transcriptional signatures described in previous studies, many of which have primarily focused on glioblastoma. To assess the similarity of our identified cell states to established transcriptional patterns, we performed gene set enrichment analysis using published signature gene sets^5,7,14,15^ (**Figure 2D**). Our neuro-, oligo-, and neuro-oligo-lineage states align with the proneural subtype^14,15^ and the neural progenitor state^5,7^. The hypoxia state strongly aligns with the mesenchymal subtype^14,15^ and state^5,7^. Our proliferative states overlap with a previously described population of cycling progenitor-like cells^7^, likely driven by the strong contribution of cell cycle genes. Interestingly, GSCs show the greatest overlap with the mesenchymal states/subtypes^5,7,15^, which include hypoxia and glycolysis-related genes. However, it is important to note that we capture hypoxia and glycolysis-related cell stress signatures in our distinct hypoxia state. This observation is likely due to the advantage that NMF offers in separating cell states that exist in a continuum through additive combinations of intrinsic features. As such, we are able to identify the specific contributions of hypoxia-associated and GSC-defining gene sets.

### Cell state composition varies across glioma tumour types

Although all cell states were shared across the three tumour types, consistent differences in cell state composition were observed (**Figure 3A, 3B and Table S7**). A notable trend was the increased proportion of tumour cells in both oligodendroglioma and astrocytoma tumours of the neuro-, oligo- or neuro-oligo-lineage cell states, 83.0% and 80.0%, respectively, compared to only 50.7% in glioblastoma tumours. Conversely, glioblastoma tumours harboured a significantly greater proportion of cells in the hypoxia, cycling and stem cell states. A larger fraction of proliferative cells is consistent with the aggressive growth of these tumours. Rapid growth outpaces the supply of a disorganised blood vessel network, creating large necrotic areas that occupy a large fraction of the total tumour volume^1^. Thus, the higher proportion of hypoxic cell states is consistent with these growth characteristics. Significant differences in the proportion of cells in a GSC state were also observed between all three tumour types (**Figure 3B**). The mean proportion of GSCs observed in oligodendrogliomas was 2.8%, 4.6% in astrocytoma and 11.4% in glioblastoma. Interestingly, this relation matches the trend in overall survival observed among the three tumour types, with a higher percentage of GSCs linked to poorer survival outcomes (**Figure S3A**). Only the proportion of GSC state was associated with survival outcomes, identifying this cell state as an important target for therapeutic development and research.

**Figure 3.**
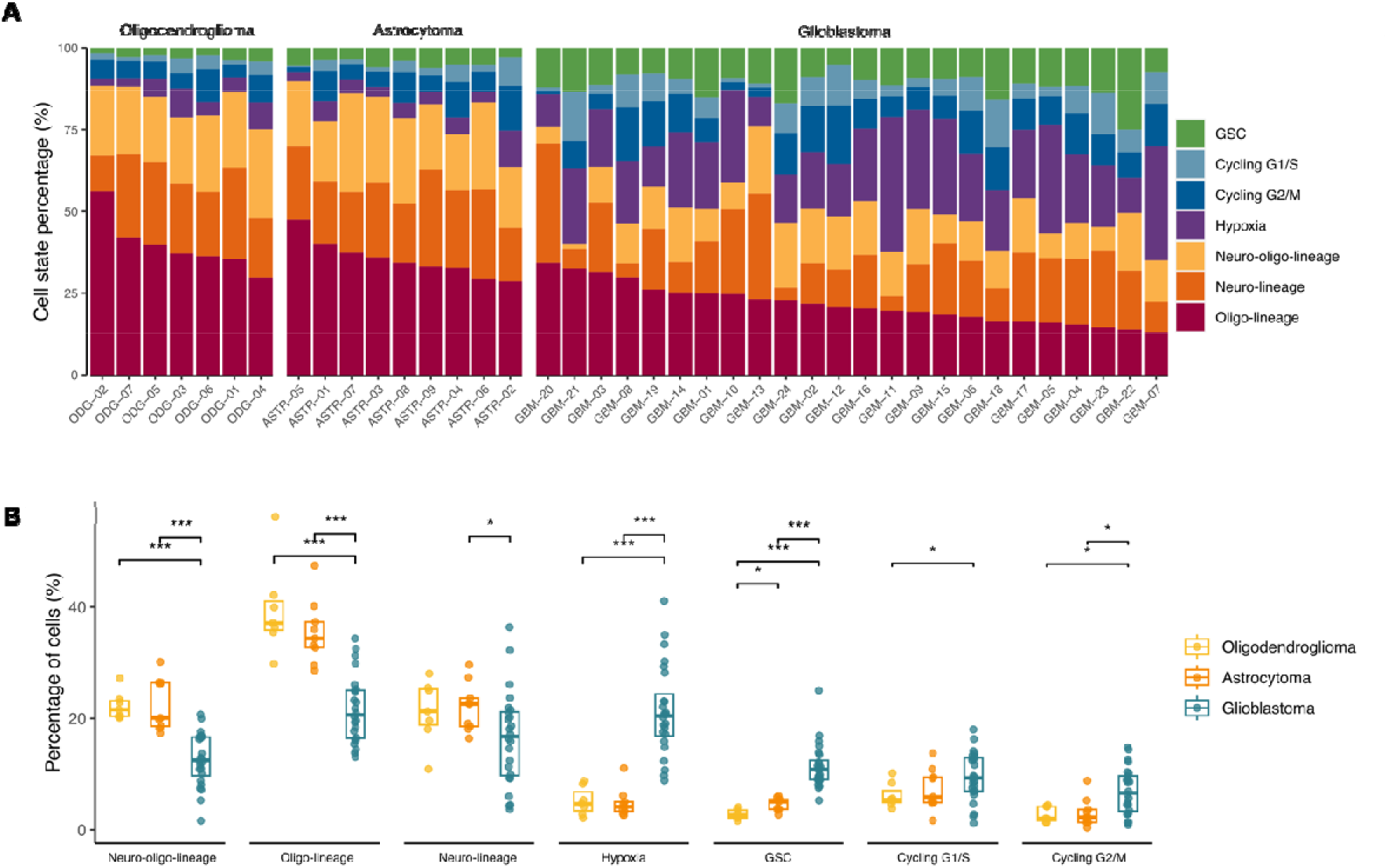
Glioma tumours comprise varying proportions of seven recurring cell states. (A) Bar plot illustrating the percentage of tumour cells in each cell state for each tumour sample. Samples are organised by tumour type: oligodendroglioma (left), astrocytoma (middle), and glioblastoma (right). (B) Box plots compare the percentage of cells in seven cell states for each tumour type. *p* values were obtained with the Wilcoxon signed-rank test. ^∗^p < 0.05; ^∗∗∗^p < 0.001.

### Tumour phylogenies identify a shared glioma stem cell population

Next, to further elucidate the nature of the GSC state population, we used clone-discriminative CNVs to reconstruct the clonal structure of each tumour sample. Distinct tumour subclones were successfully identified, containing a median number of seven, eight and nine unique CNVs for oligodendroglioma, astrocytoma and glioblastoma samples, respectively (**Figure 4A**). A CNV-based subclonal phylogeny was reconstructed for each tumour (**Figure 4B**). The number of distinct mutations separating them from the parent clone was identified for each subclone. The clonal evolution can be determined based on the acquisition of new mutations from the original parent clone. For example, focusing on the first glioblastoma patient sample (**Figure 4A**), the parent clone is identified by a gain of chromosome seven and loss of chromosomes ten and fifteen. A second clone arises with the acquisition of a subsequent deletion at chromosome thirteen, leading to the observation of two independent sub-clones. We observe greater clonal mutational diversity in glioblastomas compared with oligodendroglia and astrocytomas. These mutational patterns are consistent with those observed in clinical cohorts profiled with SNP array -based CNV profiling of bulk tumour tissue (**Figure S4A**).

**Figure 4.**
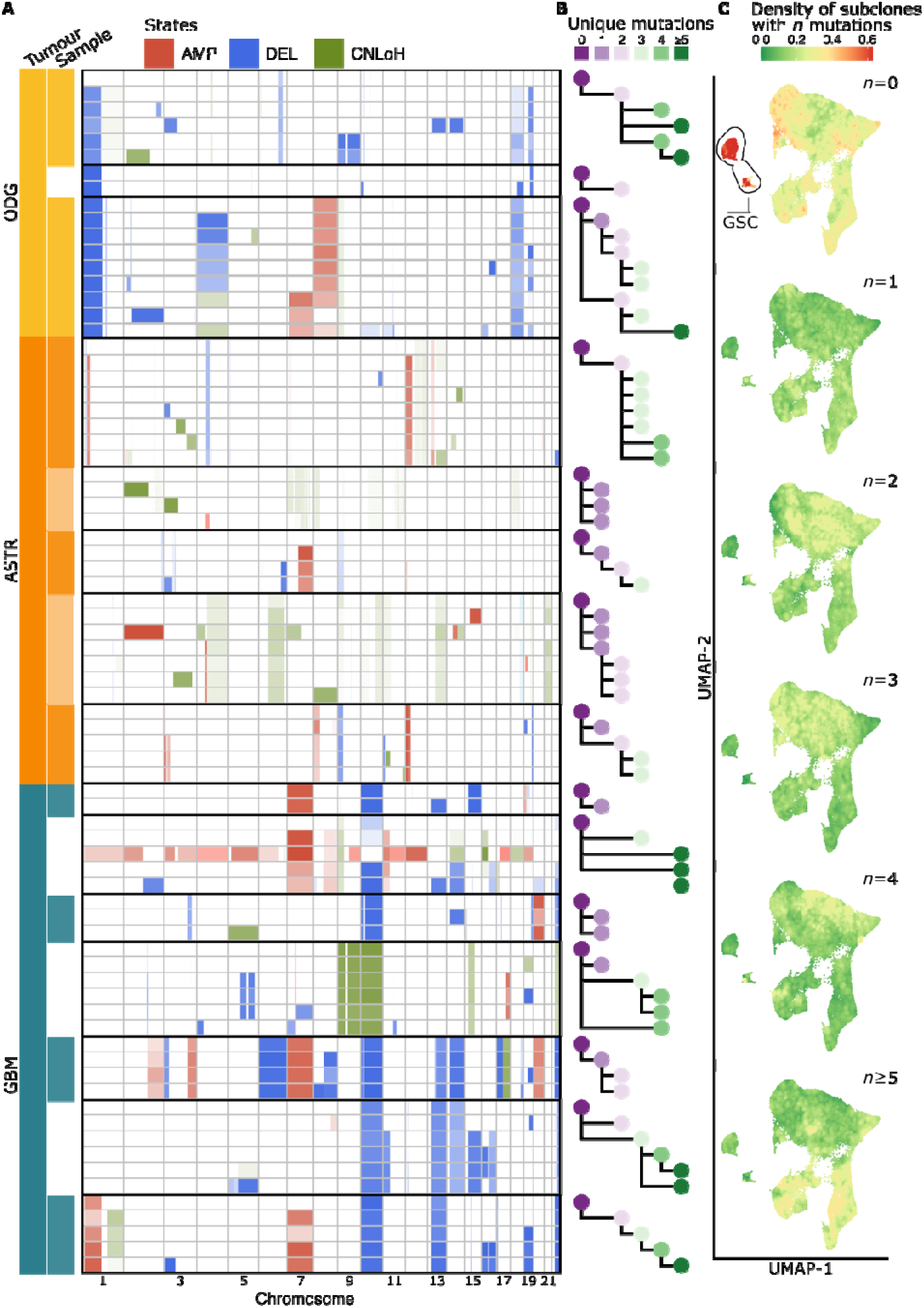
Tumour phylogenies reveal earliest tumour subclones. (A) Heatmap of tumour subclones. Rows represent individual subclones; columns represent chromosomes. CNVs that define each subclone are coloured by the type of alteration (AMP, amplification; DEL, deletion; CNLoH, copy-neutral loss of heterozygosity). (B) Tumour phylogenies for each tumour sample. Cells with the same genotype are aggregated into one node, with lines indicating the mutational history between individual subclones. Subclones are coloured by the number of distinct mutations separating them from the parent clone. (C) UMAP plots of tumour cells coloured by the density of cells assigned to subclones with *n* mutations. Red indicates an enrichment of cells from subclones with *n* mutations.

Reconstructing the clonal evolution across tumours we are able to identify cells that comprise the parent population. Using this information, we subsequently tested for the relationship between subclones and cell states. Aggregating this information across subclones over all samples, unique CNVs of the parent clone were enriched only in the population of cells corresponding to the putative GSCs (**Figure 4C and Table S8**). These results demonstrate that the GSC cell state population, across all diffuse gliomas, has a mutational signature of the parent clone. This provides further evidence that the astrocyte-like cell state identified represents diffuse glioma stem cells. In a differentiation hierarchy in which GSCs represent the apex, mutations arising in differentiating lineages are expected to be absent in the parent cells. We observe no enrichment of clonal architecture and other cell states, suggesting differentiation of GSCs does not follow a strict sequential progression.

The transcriptional similarity between GSCs and astrocytes suggested a connection with the neural stem cells (NSCs) that persist in the adult ventricular-subventricular zone, and have long been considered a potential substrate for neoplastic tansformation^16^. To explore this potential connection, we projected a pseudotime trajectory of normal mouse NSC differentiation onto our glioma dataset (**Figure S5**). The quiescent NSC population aligned with our astrocyte-like GSC population, supporting the hypothesis that this population represents a primitive stem-like state in gliomas. These observations align with recent findings that dormant glioblastoma tumour cells resembling astrocytes progress though active and differentiated stages, mirroring the structured cellular hierarchy seen in adult NSCs and their progeny^17^.

### Glioma cell states are associated with recurring genomic alterations

We hypothesised that a tumour cell’s genetic makeup may be a driver of its transcriptional phenotype. To investigate whether specific genomic alterations were enriched within distinct cell states, we used the following approach. For each genomic position, we identified cells harbouring CNVs. Using these cells, for each sample the proportion of cells in each state was calculated. We then averaged these proportions across all samples to determine an overall mean proportion per cell state. To detect significant associations, we repeated this process after randomly shuffling the cell state labels within each sample (while maintaining the overall proportion of each state) (**Table S9**).

The results of this analysis revealed several interesting trends. Cells in the neuro-, oligo- and neuro-oligo states showed a genome-wide depletion of copy number alterations (**Figure 5A-C**). These observations are consistent with the reduced exposure to mitotic errors and replication stress expected in terminally differentiated cells. Conversely, proliferating and hypoxia-associated cells showed a general enrichment of alterations (**Figure 5D,F,G**). The genomes of rapidly dividing cells are subject to a higher risk of replication-associated errors, which may result in the acquisition of alterations that confer a growth advantage or survival benefit, accumulating in descendant cell populations with subsequent divisions. Interestingly, cells in the hypoxia state were associated with significant enrichment of alterations to chromosomes seven, twelve, thirteen, fourteen, and sixteen (**Figure 5D**). Hypoxia has been demonstrated to fuel homologous recombination deficiency in many cancers^18,19^. The resulting increase in genomic instability may facilitate the acquisition of adaptive copy number changes, allowing tumour cells to overcome the microenvironmental stresses of hypoxia. These results identify a significant relationship between specific genomic alterations and distinct cell states. This suggests that a tumour cell’s genetic composition may shape its transcriptional phenotype.

**Figure 5.**
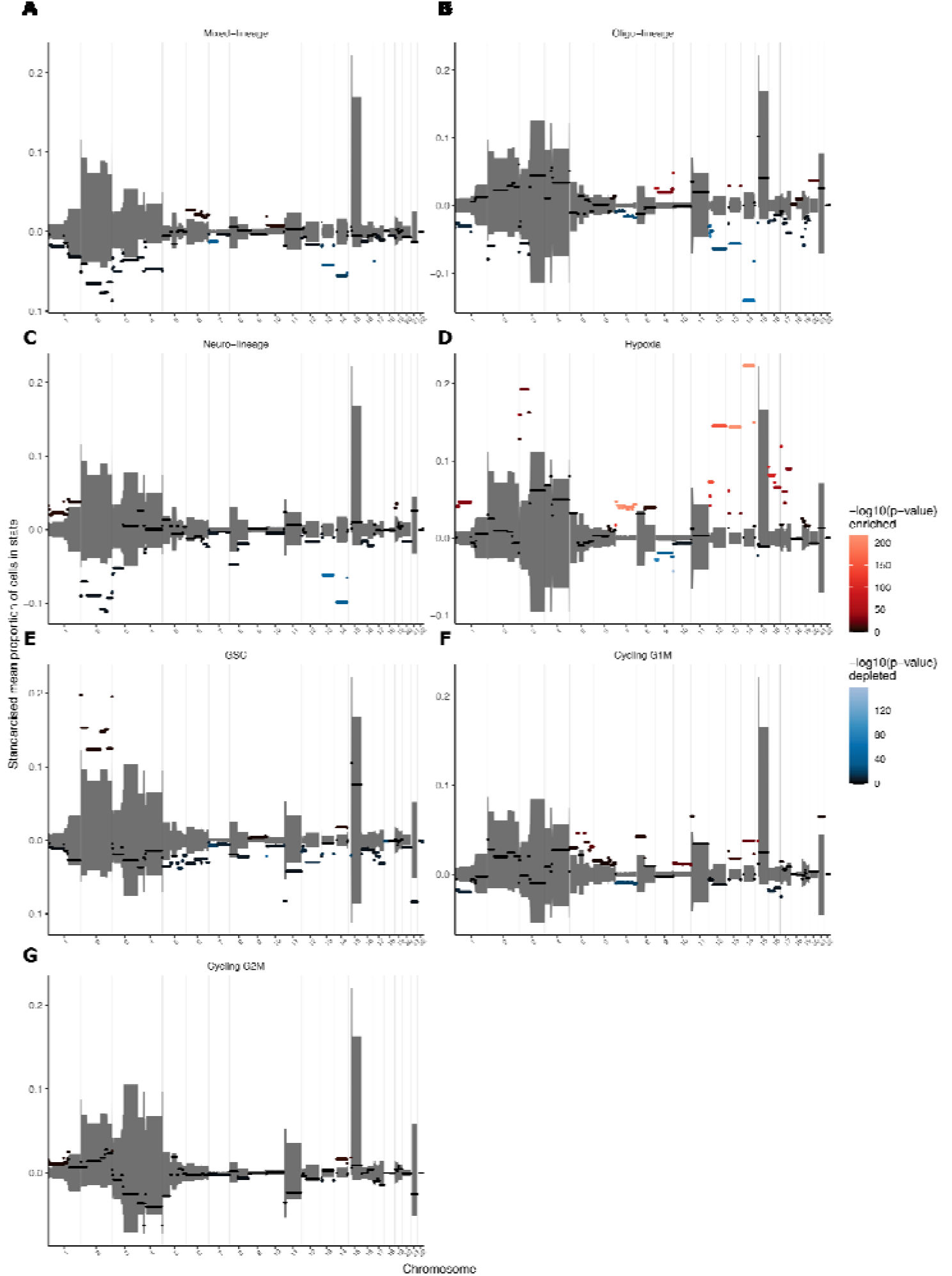
Tumour cell states are associated with recurring copy number alterations. (A-G) Panels present results for each tumour cell state, illustrating the enrichment or depletion of genetic events at each genomic position. X-axis values represent the genomic position in 1-megabase intervals. Y-axis values represent the mean proportion of cells in a cell state standardised by subtracting the mean of shuffled data. Values above zero indicate enrichment, while values below zero indicate depletion of alterations at a given genomic position. Grey shading represents the 1% to 99% confidence intervals obtained from shuffling cell states within each sample. Colours indicate the statistical significance of the observed data compared to the shuffled data.

### Validation of clonal evolution using scDNA-seq and mitochondrial DNA variants

The clonal phylogenies and observed association of GSCs with parental clones rely on CNVs inferred from the scRNA-seq data. As such, we sought to validate our observation that the parental clonal population is concordant with the GSC cell state using independent approaches. To validate the inferred CNVs, we first performed single-cell DNA sequencing (scDNA-seq) on selected samples using a custom glioma panel we designed (**Table S10**). This allowed us to directly measure DNA copy numbers in individual cells, providing a definitive confirmation of the genomic alterations identified.

The scDNA-seq data confirmed the presence of CNVs inferred by scRNA-seq (**Figure 6A and 6B**). This included whole chromosome gains and losses, such as the characteristic trisomy seven and monosomy ten. Our glioma panel design included known glioma oncogenes and tumour suppressors, hence the smaller alterations inferred by scRNA-seq, such as the focal deletion of *MEIS2* (15q14) and *TP53* (17p13) could also be validated (**Figure 6B**). Alterations to chromosomes 20-22 could not be confirmed, as the panel did not include amplicons covering these chromosomes.

**Figure 6.**
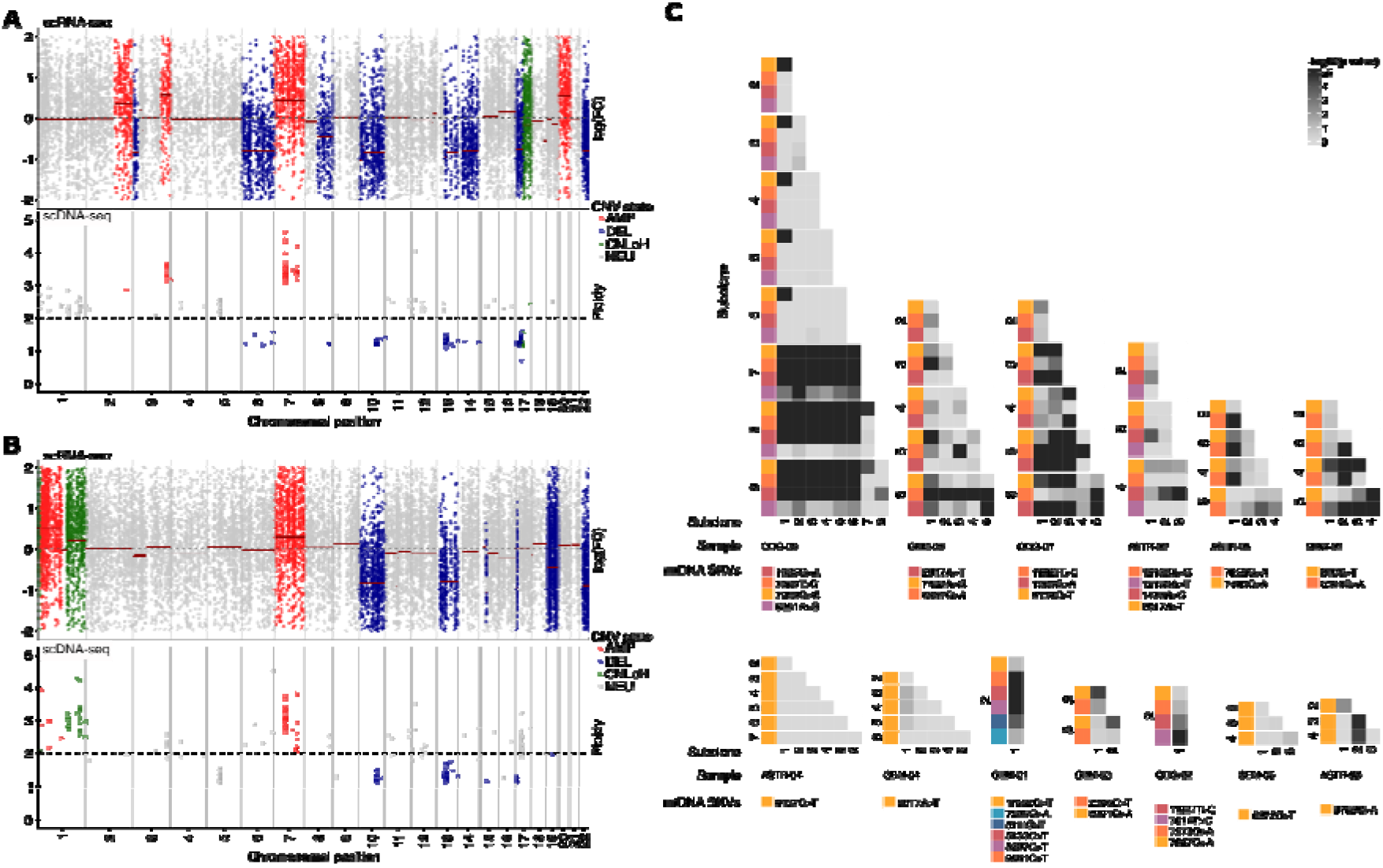
Single-cell DNA sequencing and clonally informative mitochondrial DNA (mtDNA) single nucleotide variants (SNVs) confirm copy number alterations and tumour subclones. (A) and (B) Copy number events from two tumour samples. The top panel shows the log fold-change of 100-gene window-smoothed scRNA-seq expression values. The bottom panel shows ploidy values from scDNA-seq of tissue samples from the same tumours across the same genomic regions. Colours indicate the type of genomic alteration at a given genomic position (AMP, amplification; DEL, deletion; CNLoH, copy-neutral loss of heterozygosity; NEU, neutral). (C) Heatmaps illustrate mtDNA SNV allele frequency shifts between tumour subclones. Each heatmap presents results from comparisons between subclones within an individual tumour. SNVs are demarcated by a coloured bar to the heatmap’s left and are grouped by subclone. Grey shading represents *p*-values obtained with the Wilcoxon signed-rank test, comparing SNV allele frequency value distributions between each subclone pair.

Further, to establish the accuracy of the subclones identified, we examined mitochondrial DNA (mtDNA) single nucleotide variants (SNVs). mtDNA SNVs can serve as natural genetic barcodes to recover cell lineages^20,21^. mtDNA has a higher mutation rate than nuclear DNA. As tumour subclones diverge, unique SNVs accumulate and propagate in the resulting lineages. These SNVs can then be recovered from the scRNA-seq reads. If the tumour subclones are accurate, we expect distinct mtDNA SNVs to accumulate as subclones diverge, creating significant differences in their frequencies. Testing for this, significant differences in mtDNA SNV frequencies were detected across tumour subclones (**Figure 6C**). These results demonstrate that the sub-clonal tumour architecture based on the CNVs inferred from our scRNA-seq data are accurate, lending additional support to the reconstructed phylogenies, and the relationship between the parental clone and GSC.

## Discussion

Recent single-cell analyses of adult-type diffuse gliomas have revealed malignant cells recapitulate lineages of glial differentiation. However, this work has focused on the tumour types in isolation. This has left a critical gap in our understanding of whether these transcriptional patterns are consistent and universally present across the gliomas, potentially indicating a common biological basis that could inform more effective treatment strategies. Here, we use cellular genomics combined with novel analysis methods to identify a shared cell state architecture across diffuse gliomas. We show that the glioma stem cell displays an astrocyte-like genomic signature, and confirm this observation using the somatic mutations arising from the clonal evolution of cancers.

Our findings demonstrate that despite each tumour type’s unique genetic alterations and clinical presentations, cells converge to a common, limited repertoire of seven recurring cell states. These states encompass neural lineage development (neuro-, oligo-, neuro-oligo-lineage), proliferation (cycling G1/S, cycling G2/M), cellular stress response (hypoxia), and an astrocyte-like GSC population. Overlap was observed with transcriptional patterns described in previous work within individual glioma types (**Figure 2D**). Yet, our results diverge from previous research regarding the proliferative and GSC states. Firstly, we define GSCs as having an astrocyte-like expression profile. Secondly, we find the GSC and proliferative cell states do not overlap.

Early single-cell investigations of glioma identified a gradient of stemness to differentiation within glioblastoma tumour cells. In a pioneering study, tumour cells with stem-like phenotypes were identified using a stemness signature based on genes differentially expressed between stem-like and differentiated cells derived from primary tumours^2^. This highlighted a continuous spectrum of stemness within tumours, with GSCs expressing genes associated with neural stem cell self-renewal and quiescence. Expanding on this idea by incorporating work characterising GSCs in bulk glioblastoma tumour tissue^22^, it was suggested that GSCs exist on a spectrum spanning from proneural to mesenchymal phenotypes^3^. Interestingly, consistent with the notion that stemlike cells divide at lower overall rates, GSCs defined in these studies were shown to represent a relatively quiescent founder state. Subsequent studies supported patterns of transcriptional heterogeneity from a gradient of stemness to lineage differentiation. For example, glioblastoma cells compared against GSC-enriched cells from neurospheres identified a GSC-driven tri-lineage hierarchy, with GSCs at the apex^7^. Investigations of IDH-wildtype gliomas similarly concluded astrocytomas and oligodendrogliomas share a developmental hierarchy composed of GSCs resembling neural progenitors that give rise to differentiated cells of astrocytic and oligodendrocytic lineages^4,6^. These studies share the observation that most cycling cells are GSCs.

Diverging from the concept of a unidirectional GSC-driven developmental hierarchy, it has also been suggested four primary cellular states exist in glioblastoma, and that tumour cells can reversibly transition^5^. Uniquely, the authors identify cellular proliferation within all states, albeit weighted toward oligodendrocytic and neural lineages. Summarising these developments, most findings support one of two broad concepts: a stemness gradient driven by quiescent GSCs or a neurodevelopmental hierarchy driven by proliferative GSCs. With this context we now examine our findings.

An astrocyte-like population was identified as GSCs based on their early positioning in reconstructed tumour phylogenies, supported by both scDNA-sequencing and the distribution of mtDNA SNVs. Cycling cells were identified by an analytical approach that can effectively decompose mixed transcriptional signatures to separate cellular processes. This analysis revealed proliferating cells form a distinct transcriptional group in gliomas, rather than representing a transient state blended within GSCs or the larger differentiating population. These results strongly suggest a quiescent population of astrocyte-like GSCs at the apex of a differentiation hierarchy. The non-proliferative nature of GSCs is consistent with early single-cell studies in glioblastoma^2,3^, while a differentiation hierarchy is consistent with recent pan-glioma findings^4,6,7^. Uniquely, this result repositions astrocyte-like cells as the basis of this hierarchy, despite these cells being consistently classified as belonging to the differentiated population of tumour cells. However, this converges with recent work into the glioma cell of origin. Direct genetic evidence has demonstrated that primary glioblastoma tumours arise from astrocyte-like NSCs within the human adult subventricular zone^23^. This aligns with pre-clinical evidence, in which inactivation of tumour suppressors in NSCs is necessary and sufficient for astrocytoma formation in fully penetrant mouse models^24^. In the healthy adult brain, NSCs maintain a quiescent state, sharing transcriptional similarities with parenchymal astrocytes, until activated to generate proliferating progenitors that ultimately differentiate^25^. Analogous hierarchies have been demonstrated within GSCs^26^. Indeed, a recent investigation of glioblastoma offers strong evidence for quiescent, astrocyte-like cells as GSCs, capable of establishing tumour lineages that closely resemble those of healthy adult neural stem cells^17^.

Our results suggest the shared patterns of transcriptional heterogeneity observed across the three tumour types of adult-type diffuse glioma stem from a GSC population recapitulating lineages of healthy adult NSCs. The extent to which tumour cell states resemble those of normal NSC development will likely depend on the genetic landscape of each tumour and the extent to which GSCs retain the multipotent characteristics of NSCs. While our findings identify GSCs in each tumour type, they do not reveal the specific cells of origin. The link between GSCs and the origin of these gliomas remains an unresolved question. Historically, differences in preferential localisation and molecular and clinical characteristics supported the notion that gliomagenesis occurs in the three tumour types through distinct cellular pathogenic mechanisms. Thus, the GSCs we identified may originate through distinct mechanisms in each tumour type: neoplastic transformation of existing NSCs or dedifferentiation of more lineage-restricted cells. Progress through the tumour cell hierarchy can be conceptualised by viewing cell states as stable phenotypes represented by low-energy attractor states on the proverbial Waddington landscape^27^. Different mechanisms of gliomagenesis influence GSC multipotency, determining their starting positions on this landscape^26,28^. Subsequent genetic events skew differentiation toward distinct lineages. These factors shape the unique blend of cell states observed within each tumour type, with astrocytomas and oligodendrogliomas displaying a higher proportion of terminally differentiated cells than the greater stem and cycling progenitor populations in glioblastoma. Understanding of the factors governing transitions within the tumour cell hierarchy will be crucial for designing rational treatment strategies.

One potential driver of cell state transitions is the underlying genetic instability of each tumour. Previous investigations suggest specific genomic alterations influence transcriptomic diversity but do not completely define cell states^4–8^. In this study, several notable patterns were observed. Cells in the neuro-, oligo- and neuro-oligo states showed a genome-wide depletion of copy number alterations. Conversely, proliferating and hypoxia-associated cells showed a general enrichment of alterations. Hypoxia-associated cells were specifically enriched in alterations to chromosomes seven, twelve, thirteen, fourteen, and sixteen. These findings underscore the relationship between a tumour cell’s genetic landscape and transcriptional state. Further functional validation will be required to determine the extent to which genomic alterations drive cell fate or if microenvironmental pressures create specific niches where cells with corresponding alterations outcompete others.

In summary, our study represents a significant advance in understanding the cellular complexity of adult-type diffuse gliomas. By unveiling the shared cell states across different glioma types, particularly the role of astrocyte-like GSCs, we provide a foundation for developing new therapeutic strategies that target these common elements. While our findings are compelling, they also underscore the complexity of glioma biology and the challenges in translating these insights into clinical practice. Future studies should focus on functional validation of the proposed GSC population to demonstrate self-renewal, differentiation, and enhanced tumour initiation potential. A more detailed understanding of the mechanisms underlying the formation and maintenance of the GSC state will provide insights into potential therapeutic avenues.

## Methods

### Resource availability

#### Lead contact and Materials availability

Further information and requests for resources and reagents should be directed to and will be fulfilled by Joseph Powell.

### Data and code availability

All single-cell RNA-seq data have been deposited at GEO and are publicly available as of the date of publication. All code is available from <u>github.com/powellgenomicslab/brain_cancer_paper</u> and is publicly available as of the date of publication. Any additional information required to reanalyse the data reported in this paper is available from the lead contact upon request.

### Experimental model and study participant details

#### Human tumour tissue acquisition

This study comprised adult diffuse glioma patients treated at the Prince of Wales Private Hospital in Sydney, Australia. The study was approved by Bellberry Human Research Ethics (HREC 2019-08-682), and all patients provided preoperative informed consent for participation. The study cohort comprised 26 male and 14 female patients. We included IDH mutant and wildtype gliomas, specifically astrocytomas (grades 2, 3, and 4), oligodendrogliomas (grades 2 and 3), and glioblastomas. **Table S1** summarises the clinical information.

### Method details

#### Sample preparation for single cell experiments

Freshly resected brain tumour tissue samples were collected in Gibco Hibernate-A Medium (Cat#A1247501) with neuronal cell culture supplement Gibco B-27 (Cat#17504044) and transported on ice. The tissue was rinsed with Ringer’s solution (Cat#AHF7163) to remove blood. Tissue was cut into 1-2 mm³ sections and then dissociated using the Miltenyi Biotec Brain Tumor Dissociation Kit (Cat#130-095-942), following the manufacturer’s instructions. Red blood cell lysis solution from Miltenyi Biotec (Cat#130-094-183) was used to remove the remaining red blood cells following the manufacturer’s protocol. The single-cell suspension was centrifuged three times (150 x g for 5 minutes) to remove debris. The resulting cell pellet was resuspended in B-27-supplemented Hibernate-A media and kept on ice. After passing through a 40μM filter, viability was assessed using Gibco Trypan Blue Solution (Cat#15250061). Samples with >80% viability were processed for single-cell capture using the 10x Genomics Chromium system.

#### 10x Genomics-based single-cell RNA-sequencing library preparation and sequencing

Single cells were captured for gene expression profiling using the 10x Genomics Chromium Next GEM Single Cell 3lJ kit (Cat#PN-1000130), Gem beads (Cat#PN-1000129) and partitioning oil (Cat#PN-1000129) following the manufacturer’s protocol. The cell suspension was diluted to the range of 700-1,200 cells/µL in Gibco Hibernate-A Medium (Cat#A1247501) for optimal capture efficiency, and volume was calculated for a target of 20,000 cells. cDNA was extracted from the single cell/nuclei gel bead emulsion using Dynabead clean-up mix (Cat#PN-2000048), Qiagen elution buffer (Cat#19086), reducing agent (Cat#2000087), SPRIselect reagent (Cat#B23318), and cleanup buffer (Cat#2000088). cDNA amplification was performed using Amp Mix (Cat#2000047) and a thermal cycle as follows: 98° degrees for 45 seconds, 11 cycles of (98° degrees for 20 seconds, 54° degrees for 30 seconds, 72° degrees for 20 seconds), and 72° degrees for 1 minute.

Individual libraries were pooled to target 30,000 reads per cell. Pooled libraries were sequenced at the Ramaciotti Centre for Genomics (University of New South Wales Sydney, Australia) on an Illumina NovaSeq 6000 system using NovaSeq S4 v1.5 200 cycle flowcells (Cat#20028313). Loading concentration was 294 pM with Illumina PhiX Control v3 spiked at 1%. BCL files were demultiplexed and converted to FASTQ format using the 10x Genomics Cell Ranger mkfastq pipeline. Reads were aligned to the human genome reference (GRCh38), cell and UMI barcodes were demultiplexed, and UMI counts were produced with the 10x Genomics Cell Ranger count pipeline (version 6.0.2) on a high-performance cluster with a 3.10.0-1127.el7.x86_64 operating system.

### Quantification and Statistical Analysis

#### Single-cell RNA-seq data preprocessing and quality control

Analysis of scRNA-seq data was performed with the Seurat R package version 5.0.1^29^. To identify cell-containing droplets, the CellBender version 0.2.0 remove-background tool was run for 150 epochs with a false positive rate of 0.01^30^. Droplets with a cell containment probability score of less than 0.5 were excluded from further analysis. To remove any remaining cell-free droplets, the nuclear fraction was calculated with the DropletQC R package, which quantifies the fraction of RNA originating from nuclear-unspliced pre-mRNA for each droplet^31^. The remaining cells with more than 15% of reads mapping to the mitochondrial genome were also excluded from the analysis. Next, heterotypic doublets containing cells of the four main types (tumour, myeloid, oligodendrocyte or T cells), were identified and removed based on co-expression of marker genes using the scds R package version 1.14.0^32^. Cells from all samples were integrated into a single dataset using the rliger R package (version 1.0.1)^33^, setting the number of factors (k) to 30. To visualise the integrated dataset, we applied quantile normalization and embedded it into a two-dimensional space using Uniform Manifold Approximation and Projection (UMAP) with a cosine distance metric implemented with the rliger R package. A second UMAP was also calculated using tumour cells from each sample.

### Cell-type classification of single-cell RNA-sequencing data

Cells passing quality control formed distinct clusters, the identity of which were determined by examining the expression of marker genes characteristic of each cell type: tumour cells (*PTPRZ1*, *BCAN*, *SOX4*, *EGFR*, *IGFBP2*, *TUBA1A*), myeloid cells (*C1QA*, *AIF1*, *LAPTM5*, *CD14*, *FCER1G*, *FCGR3A*, *TYROBP*, *CSF1R*), T cells (*CD2*, *CD3D*, *CD3E*, *CD3G*), and oligodendrocytes (*MOG*, *MAG*, *CNP*, *MBP*, *TF*, *PLP1*, *CLDN11*). The identity of subclusters of less common cell types were determined with the following marker genes: astrocytes (*SOX9*, *GFAP*, *AQP4*), B cells (*MS4A1*, *TCL1A*, *CD27*), dendritic cells (*CLEC4C*, *CD74*, *P2RY14*, *MZB1*), endothelial cells (*CD34*, *CLEC14A*, *VWF*, *CLDN5*), epithelial cells (*PIFO*, *CFAP276*, *RSPH1*), erythrocytes (*HBA1*, *HBA2*), fibroblasts (*COL5A1*, *COL1A1*, *COL1A2*), GABAergic neurons (*GAD1*, *GAD2*), glutamatergic neurons (*SLC17A7*, *SLC17A6*), keratinocytes (*KRT13*, *KRT6A*, *KRT5*, *KRT19*, *KRT15*, *TACSTD2*, *LY6D*), mast cells (*TPSAB1*, *TPSB2*, *KIT*, *GATA1*, *GATA2*), natural killer cells (*CCL5*, *XCL2*, *XCL1*), oligodendrocyte progenitor cells (*GPR17*, *OLIG1*, *OLIG2*, *PDGFRA*), plasmas cells (*IGLC2*, *IGHG1*, *IGKC*, *IGHG2*, *IGHG3*, *IGHGP*, *IGLC3*, *JCHAIN*) and vascular leptomeningeal cells (*ENPEP*, *FBLN1*, *ITIH5*).

### Copy-number variation analysis and tumour phylogeny reconstruction from scRNA-seq data

The Numbat package (version 1.3.2.1) was used to infer each tumour’s CNVs, clonal architecture, and evolutionary history^11^. Numbat leverages signals of allelic imbalance in expressed heterozygous SNPs and shifts in expression magnitude (reflecting chromosomal dosage) to detect underlying CNVs. Cell-level phased allele counts were generated using the pileup_and_phase.R script included in the Numbat R package. Then, leveraging the presence of non-tumour tissue in each sample, a gene expression reference was created using normal cell types (with a minimum of 30 captured cells) with the aggregate_counts function. Numbat was then run with default parameters, except for max_iter, which was increased to 20. Tumour subclones and the tumour phylogeny are inferred through an iterative process that alternates between identifying CNVs within clonal lineages and reconstructing the single-cell lineage tree based on the inferred CNV probabilities. For this cohort of samples, we found that it was necessary to increase the number of iterations to 20 for clonality predictions to stabilise. To create UMAP plots visualising tumour cell density based on the number of unique descendant mutations, the average proportion of neighbouring cells with ‘*n*’ mutations was calculated for each point. Cells in each UMAP plot were then colored according to this density, with red indicating areas enriched for subclones with ‘*n*’ mutations.

### Copy-number variation analysis of TCGA samples

Two analyses were conducted using CNV data from The Cancer Genome Atlas (TCGA) Research Network^34^. First, the mean gene-level CNV signal was calculated across oligodendroglioma, astrocytoma, and glioblastoma tumours. Second, the number of CNVs in diffuse glioma and all other TCGA tumour types was determined. For the first analysis, clinical and gene-level CNV data for all diffuse glioma TCGA samples was accessed through the TCGAbiolinks R package (version 2.28.4)^35^. Diagnoses were reclassified for consistency with the fifth edition of the World Health Organisation Classification of Tumors of the Central Nervous System (WHO CNS5) guidelines^1^, using a recent reclassification incorporating relevant molecular features^36^. A gene-level mean CNV value was then calculated separately for each tumour type. For the second analysis, segment-level CNV data was accessed through TCGAbiolinks and glioma samples were reclassified as described above. Non-tumour samples and segments smaller than 1Mb in size were excluded, and the whole genome doubling status was determined from a recent study characterising CNV patterns in human cancer^37^.

### Cell state delineation with non-negative matrix factorisation

Non-negative matrix factorisation (NMF) was employed to extract significant features from the cell-by-gene expression matrix, consisting of tumour cells from all glioma samples, as implemented in the RcppML R package (version 0.3.7)^38^. The factorisation was performed with default parameters, except for a tolerance of 1x10^-8^ and a maximum of 1,000 iterations. For each rank parameter (from 2 to 25), the factorization was repeated 40 times, each initialised with a random seed. Two measures were used to select the optimal rank parameter: mean squared error (MSE) and normalised mutual information (NMI). MSE measures the average squared difference between the original data and the approximation created by the matrix factorisation. A lower MSE indicates a better fit. NMI measures the reduction in the entropy of class labels (in this case, cells assigned to a factor) obtained if the cluster labels are known. In this case, the labels are unknown, but an increase in NMI values suggests more stable clusters. An NMI of 1 would mean a perfect clustering agreement. Considering both MSE and NMI, a rank parameter of seven provided the best balance between producing stable patterns and accurately representing the data.

### Survival analysis of TCGA glioma samples

Adult-type diffuse glioma patient clinical information generated by TCGA Research Network was obtained using the RTCGA.clinical R package (version 20151101.30.0)^39^. Guided by a recent study incorporating molecular features^36^, tumours were reclassified to align with WHO CNS5 guidelines. Kaplan-Meier survival curves were generated based on patient vital status, days to last follow-up, and days to death, with the survival R package (version 3.5.5)^40^.

### Cell state characterisation with gene set enrichment analysis

To explore the biological functions associated with each of the seven cell states, gene set enrichment analysis (GSEA) was conducted using the fgsea R package (version 1.26.0)^41^. GSEA identifies gene sets that are significantly enriched at the top or bottom of a ranked gene list. This ranking was created using the gene-by-factor matrix from the NMF analysis. Hallmark and ontology gene sets, representing well-defined biological states, processes, and cellular components, were sourced from the Molecular Signatures Database^42^. Published subtype gene signatures obtained from bulk gene expression profiling^14^, and cell state signatures generated through single-cell gene expression profiling^3,5,7,15^, were obtained from the supplementary material of the associated publications. Finally, the Benjamini-Hochberg procedure was used to adjust the GSEA p-values to account for multiple testing.

### Statistical comparison of cell state proportions

Box plots were used to visualise differences in the proportion of cells in each identified seven cell states amongst the three tumour types: oligodendroglioma, astrocytoma, and glioblastoma. The Wilcoxon signed-rank test was used to determine whether differences in cell state proportions between tumours were statistically significant.

### Comparison with healthy adult mouse neural stem cell lineage

To investigate the transcriptional similarities between human glioma tumour cells and healthy adult mouse ventricular-subventricular zone neural stem cell (NSC) lineage, the following comparative analysis was performed. A suitable NSC lineage reference dataset was constructed by incorporating data from four studies^43–46^ using the Seurat R package version 5.0.1 integration pipeline^29^. Next, single cell trajectory analysis was performed with the Monocle 3 R package version 1.3.7^47^, producing “pseudotime” values that describe the progress of individual cells along the neurogenic lineage. The resulting pseudotime values were transferred to the query glioma dataset using the Seurat TransferData function^48^.

### Statistical assessment of genomic alteration enrichment within cell states

The following analysis was performed to determine if there is a significant association between tumour cell states and copy number changes in any genomic region. For each tumour cell, the following information was provided as input: cell state, associated CNVs (chromosome, start and end coordinates), tumour type, and patient ID. First, chromosomes were tiled with 1 Mb wide segments, and the overlap between each CNV and genome tile was determined. The proportion of cells in each state was calculated for each tile and sample, followed by the mean proportion across all samples. This process was repeated 10,000 times to create a null distribution to test for statistical significance while randomly shuffling cell states within each sample. A *p*-value was calculated for each genomic position to assess whether the observed mean was greater or lesser than expected by chance. To facilitate visualization of the results, the data was standardised by subtracting the mean of the shuffled data from the 1%/99% confidence intervals and the observed data. The results were displayed by visualising confidence intervals with grey shading and colouring log10 *p*-values by the statistical significance of the observed data compared to the shuffled data for each genomic position.

### Validation of copy number variations with single-cell DNA sequencing

Nuclei were extracted from frozen tissue for single-cell DNA sequencing via the following method. Frozen tissue pieces were first washed with a nuclei suspension buffer (NSB) solution consisting of 1xPBS, 1% BSA, and 0.1U/µL RNase inhibitor. Tissue was minced manually using a scalpel blade and incubated on ice in Sigma Aldrich EZ lysis buffer (Cat#NUC101-1KT) for 5 minutes for cellular membrane disruption. The resulting solution was then passed through a 70µm filter and centrifuged at 500xg for 5 minutes, resuspended in lysis buffer and passed through a 40µm filter before centrifugation again at 500xg for 5 minutes. The nuclei pellet was resuspended in cell buffer at a concentration of 3,500 nuclei/µL and loading volume of 35µL for capture with the Mission Bio Tapestri platform. Nuclei were loaded into a Tapestri microfluidics cartridge, encapsulated, lysed, and barcoded following the manufacturer’s protocol. Barcoded samples were then subjected to targeted PCR amplification using a custom 221-amplicon panel (**supplementary materials**), designed to cover recurring genomic alterations in diffuse glioma. Libraries were pooled for sequencing on an Illumina NextSeq 500 platform. FASTQ files were aligned to the human reference genome (GRCh37) using the BWA-MEM algorithm^49,50^, and genotyped following Genome Analysis Toolkit Best Practices^51,52^, as implemented in the MissionBio Tapestri DNA analysis pipeline (version 2.0.2). The resulting HDF5 output file was imported for further analysis in R using the rhdf5 R package (version 2.44.0). The cell-by-amplicon read counts were normalised with the following procedure. High-quality cell barcodes were defined as those with total read counts greater than one-tenth of the total read counts of the tenth-highest barcode. Each read count was then divided by the mean read count for its barcode and then divided by the median read count of the high-quality barcodes for its amplicon. Finally, ploidy values were computed by dividing the normalised counts by the median counts of the diploid (non-tumour) cells and multiplying by two.

### Subclone assessment using mitochondrial DNA single nucleotide variants

Mitochondrial DNA single nucleotide variants were employed as lineage markers to validate the subclonal structures identified from the inferred CNVs. For each sample, a separate BAM file containing reads aligned to the mitochondrial genome was created specifically for tumour cells. The tool cellsnp-lite (version 1.2.2) was then employed to call variants within the mitochondrial genome^53^. To distinguish informative variants from background noise, MQuad was executed (with the AD and DP sparse matrices output by cellsnp-lite) with default parameters except for a minimum read depth of 3^20^. For validated variants, allele frequency values were calculated for each cell. Finally, the Mann-Whitney U test (stats R package, version 4.3.1) was used to assess significant differences in allele frequencies between tumour subclones.

## Acknowledgements

We gratefully acknowledge the patients who contributed their tumour tissue, making this study possible. We extend our appreciation to the members of the Garvan Institute of Medical Research Cellular Genomics Platform for their contributions to the processing of samples for single-cell capture in this study, with particular thanks to Hira Saeed, Eric Lam, Naushad Moti, and Dominik Kaczorowski for their assistance in troubleshooting. We thank Anne Senabouth for her assistance in optimising and troubleshooting the computational analysis of scRNA-seq data. A grant from the Charlie Teo Foundation supported this work. J.E.P is supported by the National Health and Medical Research Council Fellowship (APP1107599) and with the support of the Fok Family in memory of Dr. and Mrs. Wing Kan Fok. This work was supported by a National Health and Medical Research Council project grant (APP1143163) and an Australian Research Council Discovery project (DP180101405).

## Supplementary Tables

**Supplementary Table 1** | Clinical cohort information. This file describes the clinical characteristics of the forty tumour samples captured for scRNA-seq. Each row describes a patient sample. The six columns contain the following information. Patient ID: a unique identifier for each patient. Tumour type: adult-type diffuse glioma classification (oligodendroglioma, astrocytoma, or glioblastoma). Grade: WHO CNS 5 tumour grade (1 to 4). Sex: patient’s biological sex (M: Male, F: Female). Primary/Recurrent: indicates whether the tumour sample was from the initial diagnosis or a recurrence. IDH mutation status: presence or absence of an IDH gene mutation within the tumour.

**Supplementary Table 2** | Cell type markers. This file lists the genes used for cell type classification. The first column contains the cell type, and the second column lists the corresponding gene marker.

**Supplementary Table 3** | Cell annotation metadata. This file provides cell type and state annotations for each of the 543,088 cells profiled with scRNA-seq. Each row represents an individual cell with the following information. Patient ID: a unique identifier for each patient. Tumour type: adult-type diffuse glioma classification (oligodendroglioma, astrocytoma, or glioblastoma). Cell barcode: a unique sixteen-nucleotide identifier for each cell. Note that while cell barcodes are unique within a sample, some are shared across samples. Cell type: the assigned cell type. Cell state: one of seven tumour cell states (neuro-lineage, oligo-lineage, neuro-oligo-lineage, cycling G1/S, cycling G2/M, hypoxia, or glioma stem cell).

**Supplementary Table 4** | Percentage of cell types per sample. This file contains the percentage of cell types comprising the forty tumours profiled with scRNA-seq. The four columns contain the following information. Tumour type: adult-type diffuse glioma classification (oligodendroglioma, astrocytoma, or glioblastoma). Patient ID: a unique identifier for each patient. Cell type: the assigned cell type. Percentage: the percentage of each cell type. The data in this file is associated with

**Supplementary Table 5** | Cell state markers. This file contains gene scores used for cell state classification. The first column contains the gene name. The remaining seven columns contain scores representing each gene’s contribution to a specific cell state. Higher scores indicate a stronger association between the gene and that state. Values represent the gene-by-factor output from non-negative matrix factorisation analysis. Column scores sum to one, indicating the relative weight of each gene within a cell state.

**Supplementary Table 6** | Cell state scores. This file contains cell state scores for the 317,605 tumour cells profiled with scRNA-seq. Each row represents an individual cell with the following information. Tumour type: adult-type diffuse glioma classification (oligodendroglioma, astrocytoma, or glioblastoma). Patient ID: a unique identifier for each patient. Cell barcode: A unique sixteen-nucleotide identifier for each cell. Note that while cell barcodes are unique within a sample, some are shared across samples. The remaining seven columns contain scores representing the contribution of each cell state to the transcriptional profile of each cell. Column scores sum to one. Values represent the cell-by-factor output from non-negative matrix factorisation analysis.

**Supplementary Table 7** | Percentage of cell states per sample. This file contains the percentage of tumour cells assigned to each of seven tumour cell states for each sample. The four columns contain the following information. Tumour type: adult-type diffuse glioma classification (oligodendroglioma, astrocytoma, or glioblastoma). Patient ID: a unique identifier for each patient. Cell state: one of seven tumour cell states (neuro-lineage, oligo-lineage, neuro-oligo-lineage, cycling G1/S, cycling G2/M, hypoxia, or glioma stem cell). Percentage: the percentage of tumour cells assigned to that state. The data in this file is associated with **Figure 3**.

**Supplementary Table 8** | Subclone and cell state relationships. This file provides the data used to analyse the association between tumour subclones and cell states. Each row represents a single cell with the following information. Tumour type: adult-type diffuse glioma classification (oligodendroglioma, astrocytoma, or glioblastoma). Patient ID: a unique patient identifier. Cell barcode: a unique sixteen-nucleotide identifier for each cell. Note, while cell barcodes are unique within a sample, some are shared across samples. The remaining six columns indicate the density of cells belonging to subclones with *n* (zero to five) mutations distinct from each tumour’s parental subclone. The data in this file is associated with **Figure 4**.

**Supplementary Table 9** | Enrichment of recurring copy number alterations within cell states. This file contains data used to investigate the association between copy number alterations and different cell states. The nine columns contain the following information. Cell state: one of seven tumour cell states (neuro-lineage, oligo-lineage, neuro-oligo-lineage, cycling G1/S, cycling G2/M, hypoxia, or glioma stem cell). Chromosome: GRCh38 chromosome name. Genome tile: A 1-megabase chromosomal region. Observed data: the mean proportion of cells in a given cell state exhibiting a copy number alteration. The next three columns describe the null distribution (1%, 50%, 99% quantiles), obtained by shuffling cell states 10,000 times. The remaining two columns indicate whether the observed proportion is significantly higher or lower than expected by chance (compared to the null distribution). The data in this file is associated with **Figure 5**.

**Supplementary Table 10** | Custom glioma DNA panel. This file describes a custom 221-amplicon panel designed to cover recurring genomic alterations in diffuse glioma. The four columns contain the following information about each amplicon. Chromosome: GRCh37 chromosome name. Start coordinate: GRCh37 start coordinate. End coordinate: GRCh37 end coordinate. Amplicon ID: a unique amplicon identifier.

## Supplementary Figures

**Figure S1.**
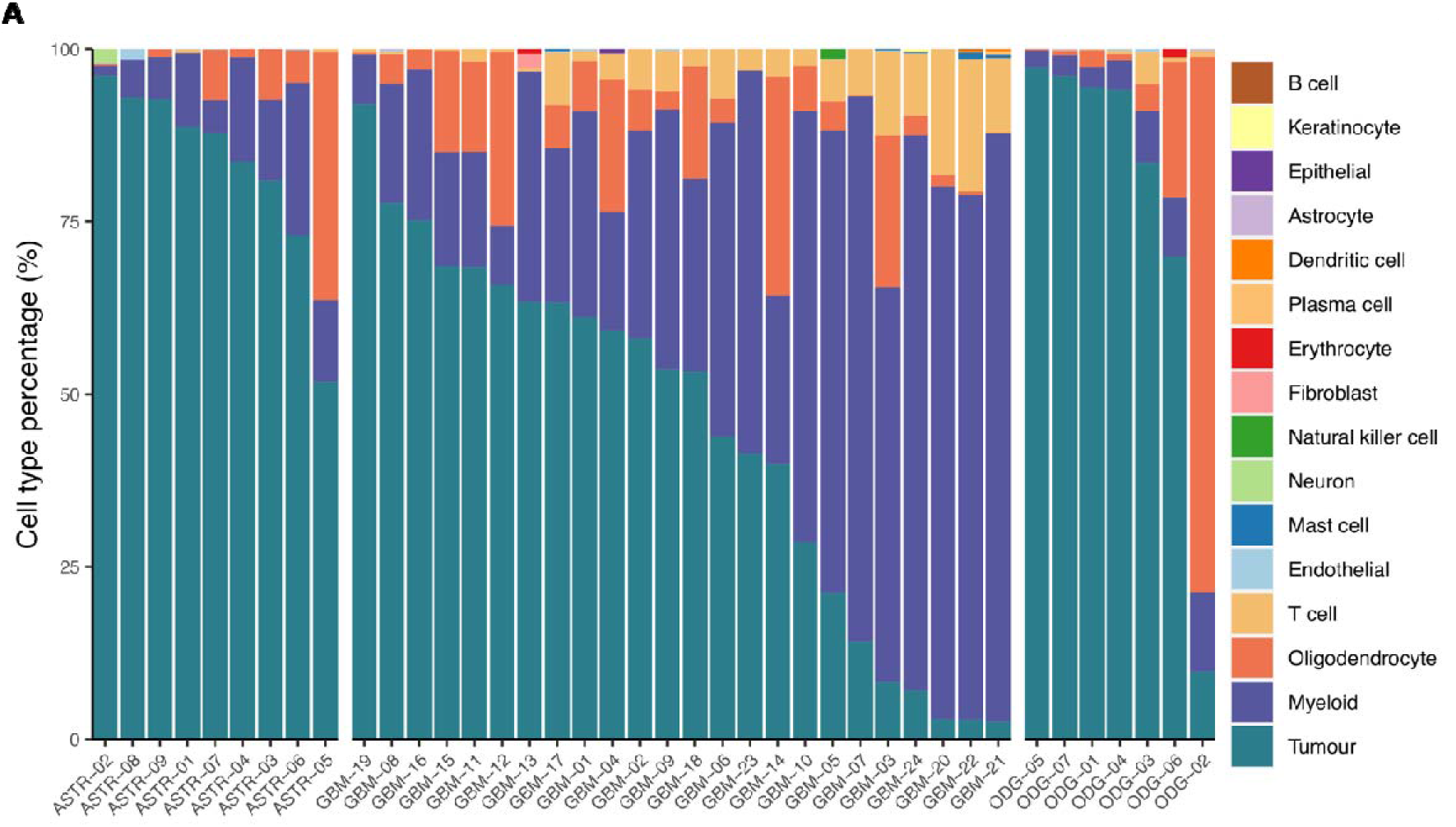
Extended classification of single cells from 40 adult-type diffuse gliomas. (A) Detailed cell type classification. The percentage of cell types that comprise each tumour is shown. The plot is split left to right into the three tumour types; astrocytoma, glioblastoma and oligodendroglioma.

**Figure S2.**
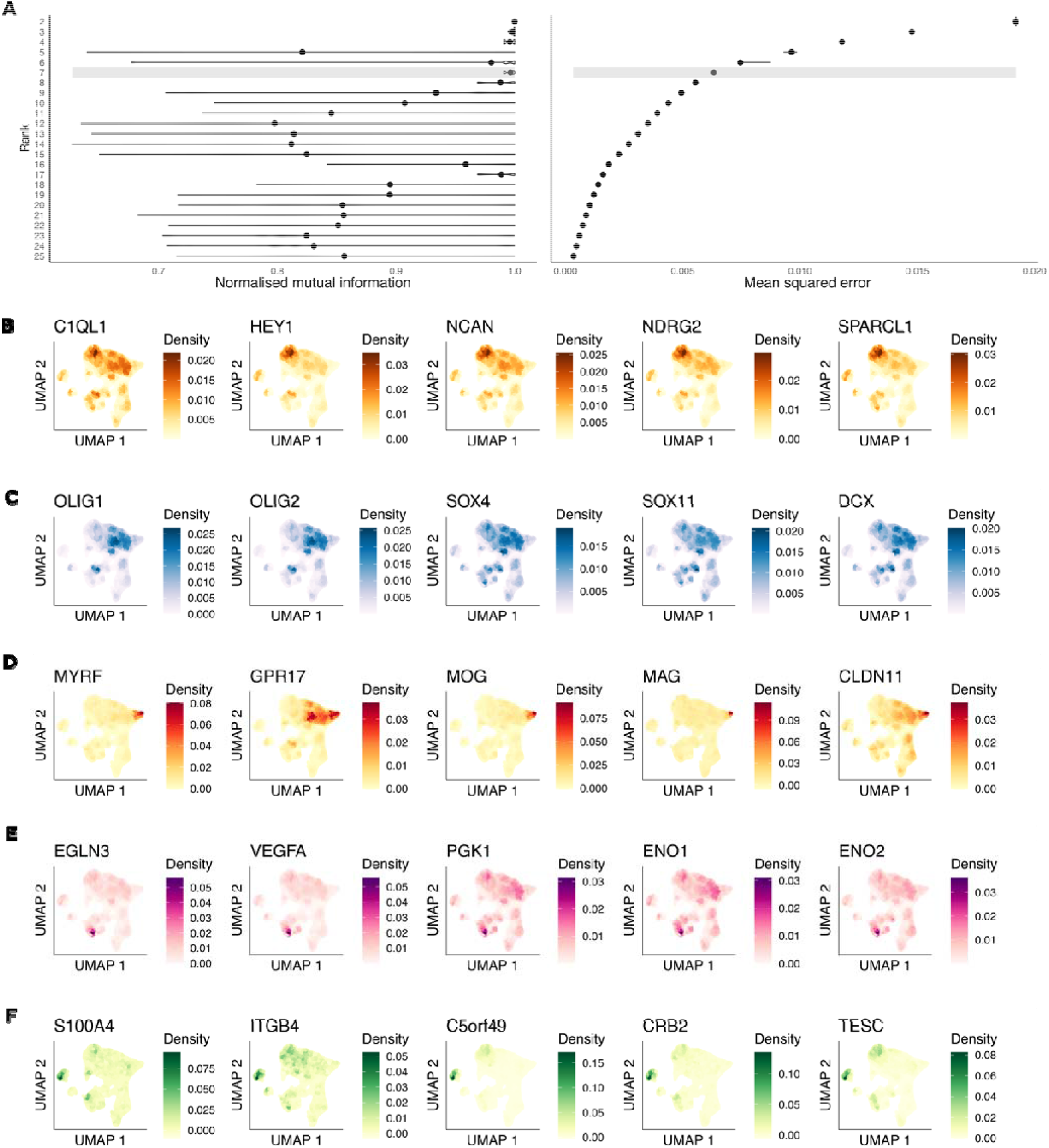
Defining an optimal NMF rank parameter and gene markers that define tumour cell states. (A) Two measures to select the optimal rank parameter: normalised mutual information (NMI) and mean squared error (MSE). NMI measures the reduction in the entropy of class labels obtained if the cluster labels are known. An increase in NMI values suggests more stable clusters. MSE measures the average squared difference between the original data and the approximation created by the matrix factorisation. A lower MSE indicates a better fit. A rank parameter of seven provided the best balance between producing stable patterns and accurately representing the data. Additional gene markers characteristic of the neuro-lineage (B), neuro-oligo-lineage (C), oligo-lineage (D), hypoxia (E) and glioma stem cell (F) states are visualised on UMAP embedding plots of tumour cells from all samples.

**Figure S3.**
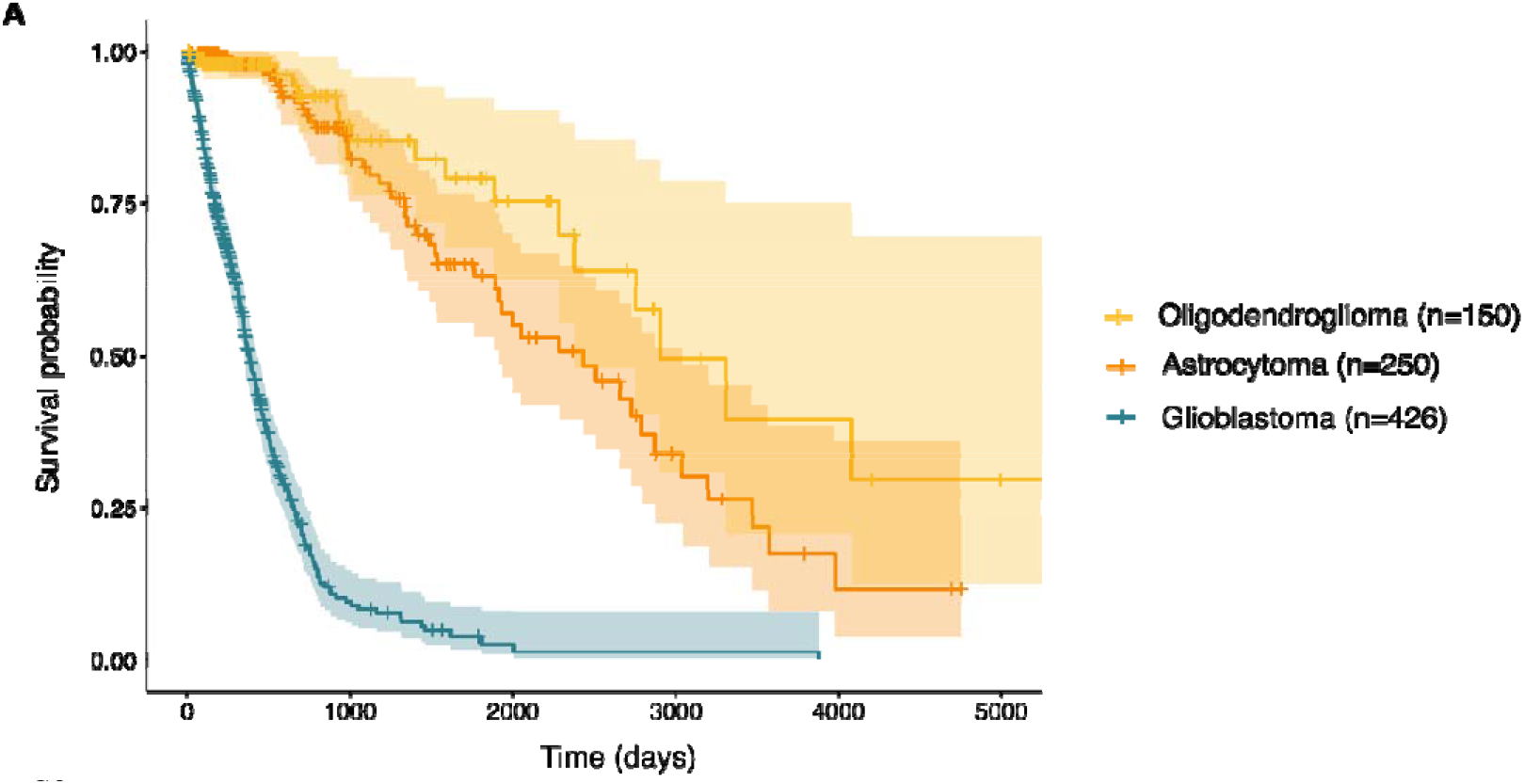
Kaplan-Meier curves demonstrating overall survival of 826 Cancer Genome Atlas project adult-type diffuse glioma patients reclassified to be consistent with 2021 WHO Classification of Tumors of the Central Nervous System guidelines.

**Figure S4.**
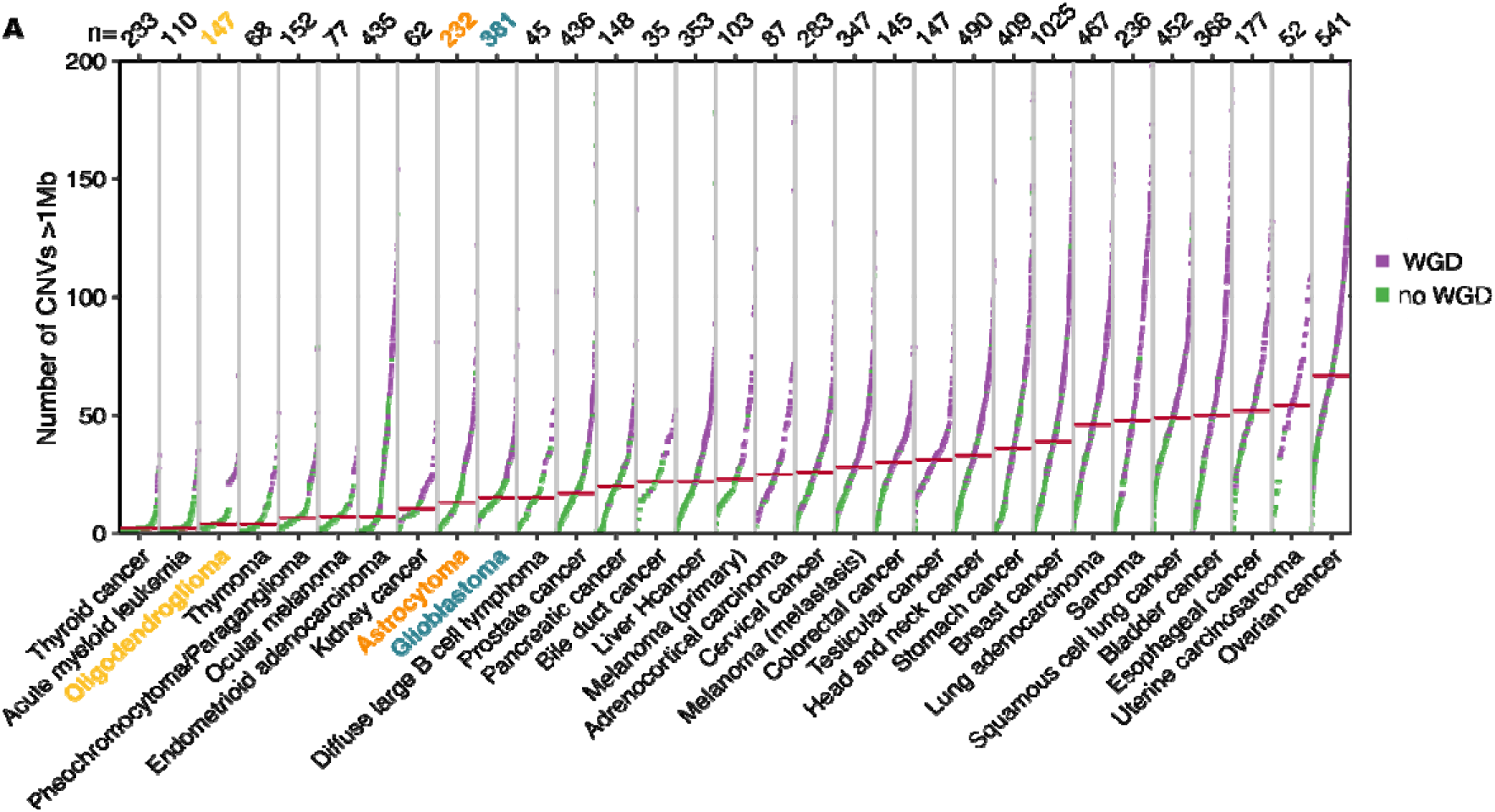
Number of CNVs of all Cancer Genome Atlas project samples split by tumour type. Colours represent the genome doubling status of each sample: non-genome doubled (green) and genome-doubled (purple). Top, the number of available samples of each tumour type.

**Figure S5.**
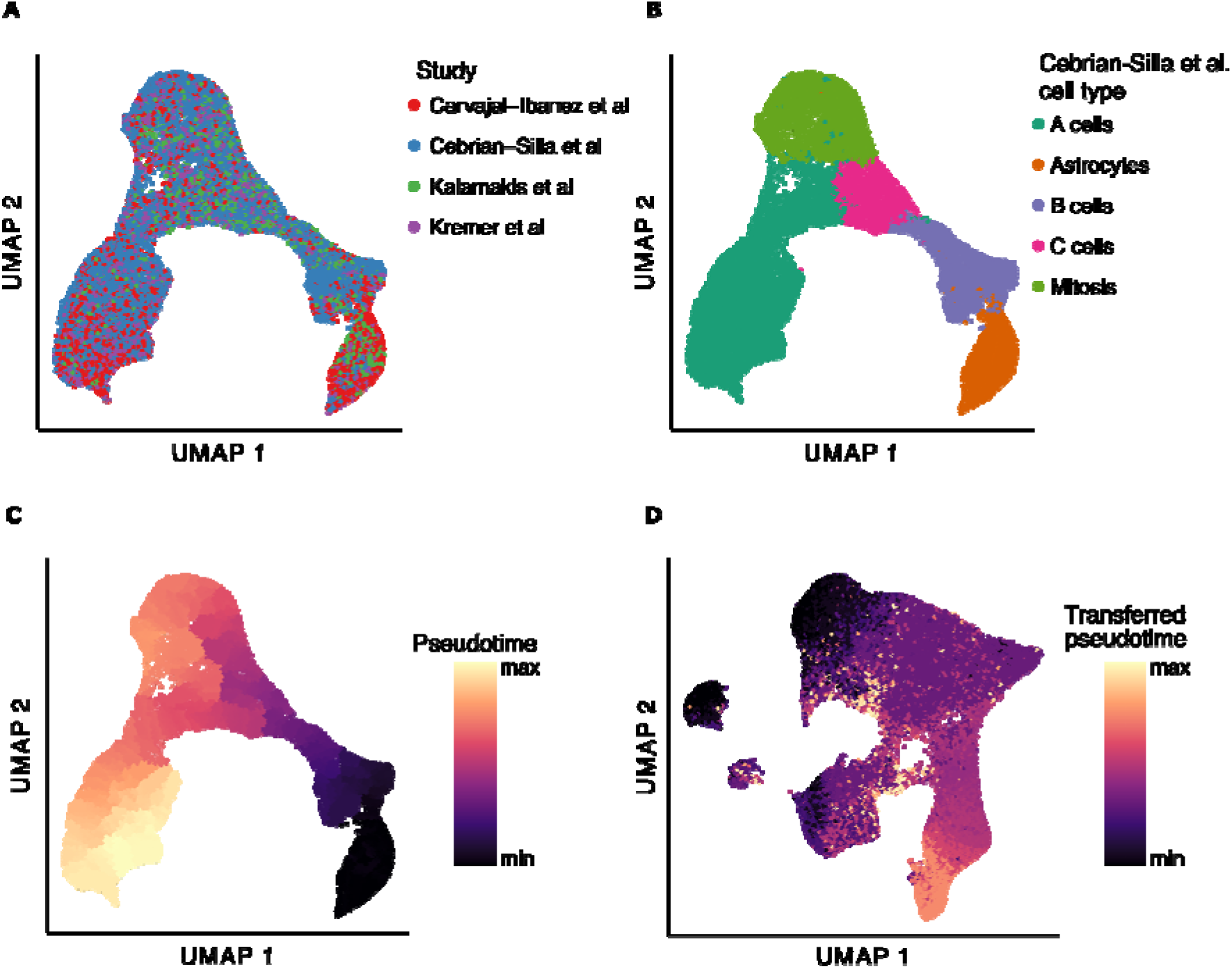
Comparison of the adult mouse ventricular-subventricular zone (V-SVZ) neural stem cell (NSC) lineage and human adult-type diffuse glioma cells. (A) UMAP embedding of the integrated reference NSC dataset with cells colored by each of the constituent four studies. (B) UMAP embedding of the reference dataset colored by cell type. (C) UMAP embedding of the reference dataset colored by pseudotime values inferred with Monocle 3. (D) UMAP embedding of the glioma dataset colored by transferred pseudotime values.

## Notes

### Competing Interest Statement

The authors have declared no competing interest.

